# Super-Beacons: open-source probes with spontaneous tuneable blinking compatible with live-cell super-resolution microscopy

**DOI:** 10.1101/2020.01.20.912311

**Authors:** Pedro M. Pereira, Nils Gustafsson, Mark Marsh, Musa M. Mhlanga, Ricardo Henriques

## Abstract

Localization based super-resolution microscopy relies on the detection of individual molecules cycling between fluorescent and non-fluorescent states. These transitions are commonly regulated by high-intensity illumination, imposing constrains to imaging hardware and producing sample photodamage. Here, we propose single-molecule self-quenching as a mechanism to generate spontaneous photoswitching independent of illumination. To demonstrate this principle, we developed a new class of DNA-based open-source Super-Resolution probes named Super-Beacons, with photoswitching kinetics that can be tuned structurally, thermally and chemically. The potential of these probes for live-cell friendly Super-Resolution Microscopy without high-illumination or toxic imaging buffers is revealed by imaging Interferon Inducible Transmembrane proteins (IFITMs) at sub-100nm resolutions.

## Introduction

Super-Resolution Microscopy (SRM) encapsulates optical imaging methods capable of bypassing the ∼250nm resolution limit imposed by diffraction (1). Their resolving power approaches that of electron microscopy (2) while keeping benefits of fluorescence imaging such as molecular specific labelling and potential for live-cell imaging (3). Single-Molecule Localization Microscopy (SMLM) is a well-establish set of SRM approaches, particularly popular due to their capacity to achieve near-molecular scale resolution (<50nm) in relatively simple imaging equipment (4, 5). SMLM-based techniques achieve this nanoscale resolution by exploiting fluorophore photoswitching (6–9) or emission fluctuation (10, 11). An SMLM acquisition aims to capture individualized fluorophores transitioning between non-emitting and emitting states. The analysis of a sequence of images containing this information then allows the creation of a Super-Resolution image where the presence and location of these fluorophores can be better-discriminated (12). To achieve a considerable resolution increase (10 fold increase when compared to diffraction-limited approaches), these methods rely on specialized labels whose switching kinetics are frequently modulated by intense illumination (13).

However, this requirement constrains these approaches to microscopes capable of high-intensity illumination and limits compatibility with live-cell imaging due to phototoxicity (3, 14, 15). Exceptions exist, such as genetically encoded fluorophores that photoswitch at low-intensity illumination, these include Dreiklang (16), Skylan (17) and SPOON (18). SMLM based on Point Accumulation for Imaging in Nanoscale Topography (PAINT) exploits an alternative mode of blinking, accomplished through transient binding of fluorescent probes to target molecules (19–21). PAINT has the benefit of not requiring high-intensity illumination, as blinking arises from binding kinetics. However, it is often limited to TIRF or Spinning-Disk microscopy, due to the presence of considerable background fluorescence from unbound freely diffusing probes (22). Recent studies have started to address this issue (23).

We propose a new class of Super-Resolution fluorescent probes for SMLM, dubbed Super-Beacons (Fig. 1). With these molecules, the transition between emitting and non-emitting states is stochastic and independent of illumination. This feature eliminates the need for photodamaging illumination or photoswitching inducing buffers (Fig. 1a). At the molecular level this is achieved through a DNA scaffold resembling Molecular-Beacons (Fig. 1b) (24), altered to achieve high-efficient photoswitching instead of oligonucleotide sensing (Fig. 1c) (25). We characterized the photoswitching properties of Super-Beacons 1d-e) and demonstrate their applicability for SMLM in live-cell compatible imaging conditions.

**Fig. 1.**
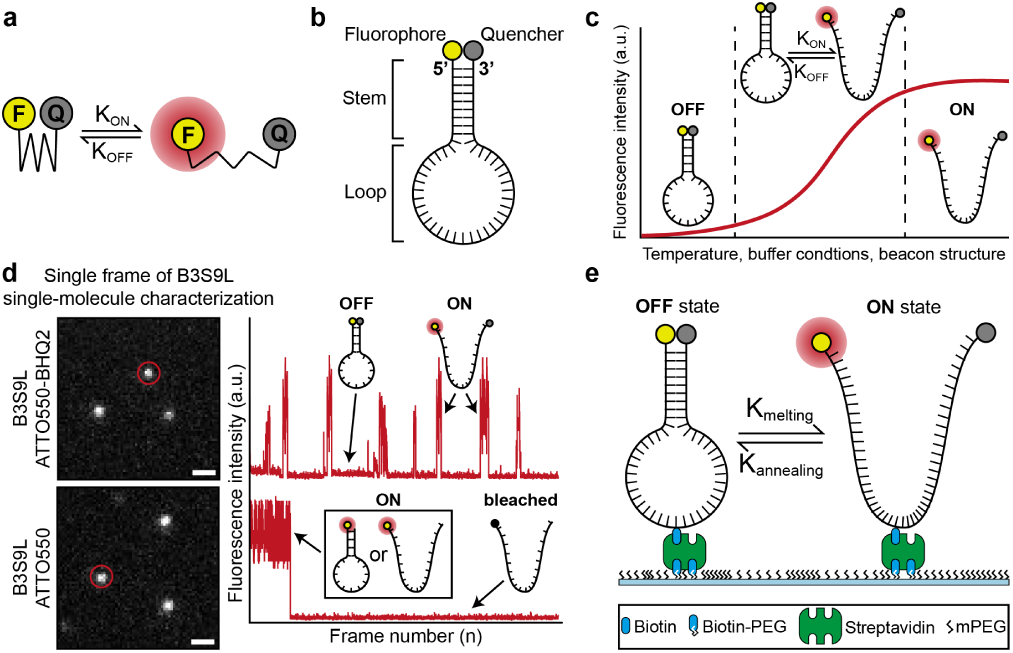
Mechanism of Super-Beacon photoswitching. a) Model of transient quenching mediated photoswitching. b) Super-Beacon structure. c) Intensity plot based on the annealing profile of Super-Beacons in different conditions. d) (left) Single-molecule *in vitro* characterization (Scale bars are 1 µm). (right) Intensity oscillation profile of highlighted molecules, showing open and closed Super-Beacon conformation. e) Schematic representation of Super-Beacons on a coverslip surface for *in vitro* characterization by single-molecule imaging.

## Results

### Super-Beacons probe design

We hypothesized that fluorophore photoswitching can be induced by promoting transient interactions between a fluorophore-quencher pair in a molecule that exhibits two conformational states: “closed” - quenched due to fluorophore and quencher contact; “open” - fluorescent where the fluorophore and quencher are not in proximity (Fig. 1a) (26). The transition between these conformations thus influences the lifetime of emitting and non-emitting states. We designed a new class of photoswitchable probes using these general principles. We chose a DNA-hairpin scaffold due to its flexibility and easy synthesis, a structure resembling the well-characterized Molecular-Beacons (Fig. 1b) (27–30).

Our probe dubbed a Super-Beacon (SB), is a single-stranded DNA (ssDNA) oligo-nucleotide with a fluorophore and quencher covalently bound to opposing ends (Fig. 1b). Short, complementary, terminal sequences promote self-hybridization resulting in the stable formation of the hairpin shaped secondary structure, hereafter called the “closed”-state. This state, dominant in thermal equilibrium, is non-emitting due to resonant energy transfer, collision and contact with a dark quencher (31, 32). The probe can reversibly transition to a short-lived fluorescent “open” state in a stochastic manner (Fig. 1c). The rate of transition between states is modulated by temperature, chemical environment and choice of structure (oligo-nucleotide sequence) (25, 33) (Fig. 1c). To demonstrate the principles of SBs, we designed a short probe with the sequence 5’-ATTO550 ACG TTT T[Biotin-dT]T TTT CGT BHQ2-3’called B3S9L-ATTO550-BHQ2. A naming convention that relates to it being a Super-Beacon (<u>B</u>3S9L) using 3 base pairs in the Stem region (B<u>3S</u>9L) and 9 poly-T bases in the Loop (B3S<u>9L</u>), an ATTO550 fluorophore conjugated to 5’-end and a Black Hole Quencher 2 (BHQ2) conjugated to 3’-end. A control structure without quencher was designed 5’-ATTO550 ACG TTT T[Biotin-dT]T TTT CGT −3’. Both probes have an internal modification to the thymine base at the centre of the hairpin loop to biotinylated thymine. This feature allows their binding to streptavidin-conjugated labelling agents such as antibodies or treated coverslips for optical characterization (Fig. 1d,e).

### Super-Beacons *in vitro* characterization

To characterize SBs as photoswitching probes, we imaged their photophysical behaviour while immobilized on a coverslip using Total Internal Reflection Fluorescence microscopy (TIRF) (Fig. 1d,e) (25). B3S9L-ATTO550-BHQ2 SBs were linked to a coverslip surface at high dilution, to achieve sparse spatial distribution and negligible detectable molecular spatial overlap. In SMLM, it is expected that probes switch stochastically between a non-fluorescent (off) state and fluorescent (on) state. The on-state lifetime, *t*_on_, should, ideally, be similar to the acquisition rate (10s of milliseconds) and ratio of on-state to off-state lifetimes, *r* = *t*_on_*/t*_off_, at least 1/50 (34). Following the protocols described in the Methods section, we measured the switching state lifetimes of our SB structure (Fig. 2a). In these conditions, r=1/50 is achieved at 250 W.cm^−2^, demonstrating desirable photoswitching at 5-fold less illumination than for our control probe. Additionally, the molecular design of SBs favours the closed-state conformation, an advantage over organic fluorophores for SMLM, since a high proportion are observed to be in the off-state at the beginning of acquisition (Fig. 2b).

**Fig. 2.**
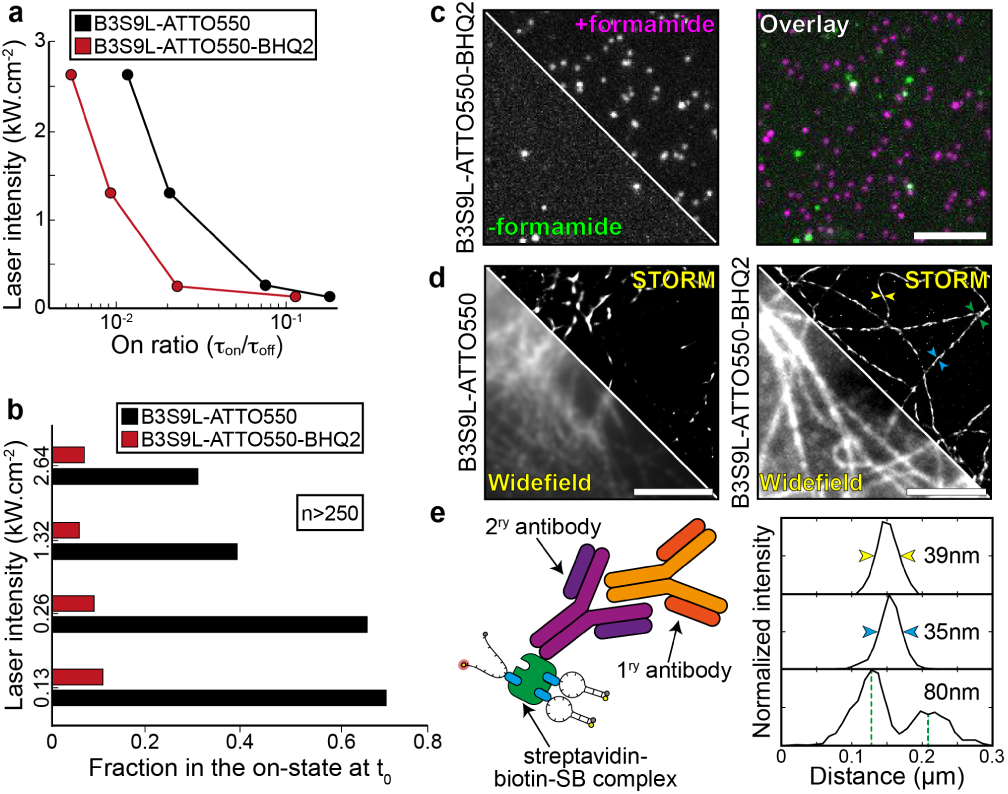
Super-Beacons characterization as SMLM probes. a) On time ratio *t*_on_/*t*_dark_ against illumination intensity, where *t*_on_ and *t*_dark_ are the weighted means of the on- and dark-state distributions respectively. (Red line) B3S9L-ATTO550-BHQ2. (Black line) B3S9L-ATTO550. b) Fraction of Super-Beacon and control probes in the on-state in the first frame of acquisition. c) Super-Beacon probe imaged in the absence (-formamide) or presence (+ formamide) of formamide and overlay of both images (overlay). d) Pre-acquisition snapshot (WF) and SMLM reconstruction (STORM) of β-tubulin immunolabelling with B3S9L SB (B3S9L-ATTO555-BHQ2) and control probe (B3S9L-ATTO555). Scale bars are 5 µm. e) Schematic representation of the strategy used to perform β-tubulin immunolabelling (left), measurement of Full With Half Maximum (FWHM) and distances between structures on the highlighted regions in d).

We then disrupted the hairpin structure with formamide, to determine if switching properties are the result of the transition between conformation states. The delayed addition of formamide and consequent melting of the hairpin structure induced an on-state indistinguishable intensity from the control structure (Fig. 2c).

### Super-Beacons as Super-Resolution probes

#### Performance as labels

To evaluate the potential of SBs as imaging probes, we took advantage of the biotin modified thymine in the SB loop to attach it to a streptavidin-conjugated antibody (Fig. 2d,e). Using these antibodies we labelled and imaged β-tubulin in fixed NIH3T3 cells (MetOH fixed as previously described (35)) with an illumination intensity of 150 W.cm^−2^ in Phosphate-Buffered Saline (PBS) buffer. These low illumination and non-toxic buffer conditions are within the acceptable range of live-cell imaging (14). Fig. 2d shows that SRM imaging with both the B3S9L-ATTO550-BHQ2 probe and B3S9L-ATTO550 control probe. SMLM analysis was performed using ThunderSTORM (36) detecting over 6×10^10^ localizations (full-frame) and generating a super-resolution reconstruction with improved contrast and resolution in comparison to the equivalent wide-field image (Fig. 2d). A low-density region of the microtubule network was selected to judge the achieved resolution, suggesting that a resolution better than 80nm, calculated by measuring the distance between two filaments (Fig. 2d,e). This experiment demonstrates that for similar low-illumination imaging conditions, using a non-toxic buffer, SBs can achieve SRM images of higher perceived quality than those obtained with the control probe (Fig. 2d).

#### Capacity for live-cell Super-Resolution Microscopy

To test the potential of SBs for live-cell SRM imaging, we used cell surface-expressed interferon-induced transmembrane (IFITM) proteins. IFITM proteins are a family of broad-spectrum inhibitors of virus replication that act primarily against enveloped viruses (37). IFITM genes have been found in numerous vertebrate species, including lampreys. In humans, IFITM-1, −2 and −3 encode proteins that are thought to act primarily, though perhaps not exclusively (38), by inhibiting viral fusion (37, 39). When expressed alone in A549 cells, IFITM-1 localises predominantly to the plasma membrane, while IFITM-2 and −3 localise preferentially in late and early endosomes, respectively (39). Thus, for human IFITM proteins at least, the distributions cover the main cellular portals through which enveloped viruses enter cells. We next investigated the potential of SBs to provide detailed information on the distribution and dynamics of C-terminally HA-tagged IFITM-1 in live-cell SMLM experiments. As IFITM-1 is primarily localized to the plasma membrane, and its C terminus is accessible on the surface of intact cells (39), we chose this protein for initial studies.

To first validate IFITM-1 SRM, we performed a fixed-cell comparison between standard STORM with AlexaFluor647 and SB-based STORM. A549 cells stably expressing HA-tagged IFITM-1 were grown on glass coverslips were fixed and labelled either with 1) an anti-HA antibody conjugated to AlexaFluor647 and subsequently embedded into a photoswitching inducing buffer (100 mM β-mercaptoethylamine); or 2) the same anti-HA antibody SB-labelled through conjugation with streptavidin (as mentioned in Fig. 2) and embedded in PBS. Both samples were then imaged through several STORM acquisitions where, for each, the illumination intensity was increased from live-cell friendly 50 W.cm^−2^ to classical STORM illumination intensities ∼2.5 kW.cm^−2^ (Fig. 3).

**Fig. 3.**
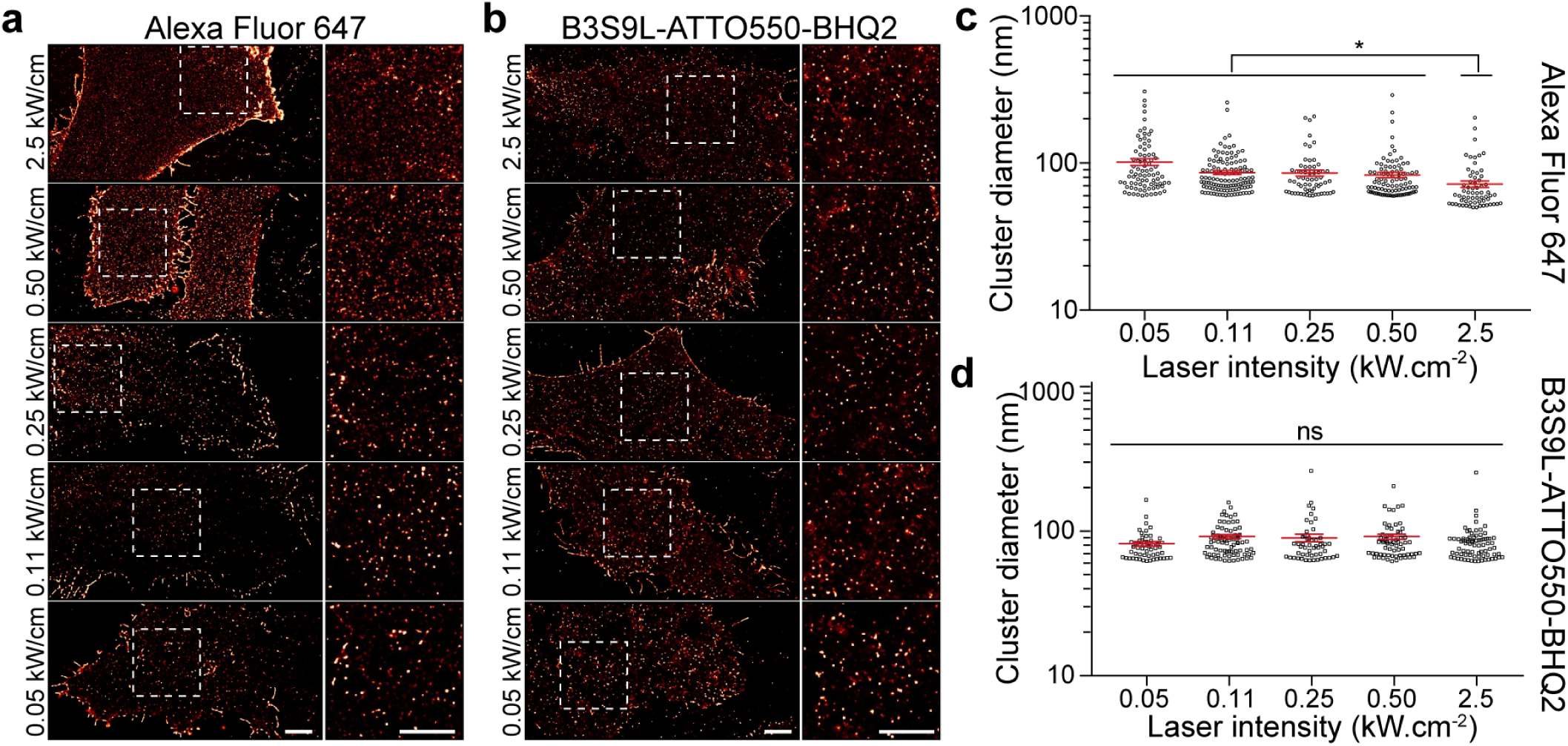
SMLM with Super-Beacons at live-cell compatible illuminations. a) IFITM1 STORM reconstruction using AlexaFluor647 or Super-Beacon B3S9L conjugated primary antibodies at increasing illumination intensities. Scale bars are 1 µm. b) SR-Tesseler cluster analysis of IFITM1 cluster diameter (nm) at increasing illumination intensities with anti-HA primary antibodies conjugated with AlexaFluor647 or Super-Beacon B3S9L.

Despite the order of magnitude difference in illumination intensity across modalities and the SB sample not containing a buffer promoting photoswitching, there were negligible visual differences in IFITM1 distribution at the cell surface, for both imaging modalities (Fig. 3a). Quantitative analysis of image quality through the SQUIRREL algorithm (40) further shows that both conditions achieve similar quality (Fig. S2). In all illumination regimes, there was a high fidelity between the SMLM reconstructions and the raw data (as seen by SQUIRREL error maps). As expected, this was not the case for AF647 that relies on photo-induced processes (Sup. Fig. S2a,b).

SMLM provides not only high-resolution structural information but also quantitative information about the subcellular distribution and molecular organization (41). Hence, we asked if the high-fidelity in protein distribution across different illumination intensity regimes was translated into the ability to extract quantitative data. To access this, we analysed IFITM1 distribution across a range of illumination intensities (from ∼ 0.05 kW *cm*^−2^ to ∼ 2.5 kW *cm*^−2^) and analysed the cluster diameter with SR-Tesseler (41). When using anti-HA-AF647 labelled antibody, we saw a significant difference (p<0.01) between the high-illumination regime (2.3 kW *cm*^−2^) and the low illumination regimes (>0.46 kW *cm*^−2^) and a corresponding increase in the mean cluster diameter (from 70 nm to values ranging between 80-100 nm). For the anti-HA-streptavidin-SB Ab conjugate, we obtained an average cluster diameter of 80 nm. Importantly, we saw no difference (p>0.1) between the different illumination regimes. The 10 nm discrepancy in size between both approaches was indicative of linker length between the fluorophore and the antibody (NHS for AF647 vs streptavidin-biotin for SB). Adjusting the illumination intensity affected the measured cluster diameter when using AlexFluor647, due to the photo-physical dependence of the switching rates, whereas there was no difference when using the SB.

Having confirmed that SMLM of HA-tagged IFITM1 is possible, live-cell imaging was performed at 0.132 kW *cm*^−2^, 561 nm illumination with exposure time 20 ms in Ringers solution (Fig. 4). To do this A549 (IFITM1-HA) were incubated with SB labelled Anti-HA primary antibody for 30 min, at 37°C in Ringers solution (cells were washed twice with Ringers solution before imaging). There was a small deterioration in the localization precision compared to fixed cell imaging, *σ*_*loc*_=30.7 nm, which is expected due to the reduced photon count when using shorter exposure times and the motion blurring of the molecule during the exposure. The dynamics of the IFITM1 protein could be visualized by rendering of the localizations with a lookup table indicating time (Fig. 4a). The increase in the spatial extent of these clusters in comparison to fixed cells was too large to be accounted for by the decrease in localization precision which could be suggestive of diffusion of the molecule within confined regions. Thin trails of localizations between these regions were also seen in few instances.

**Fig. 4.**
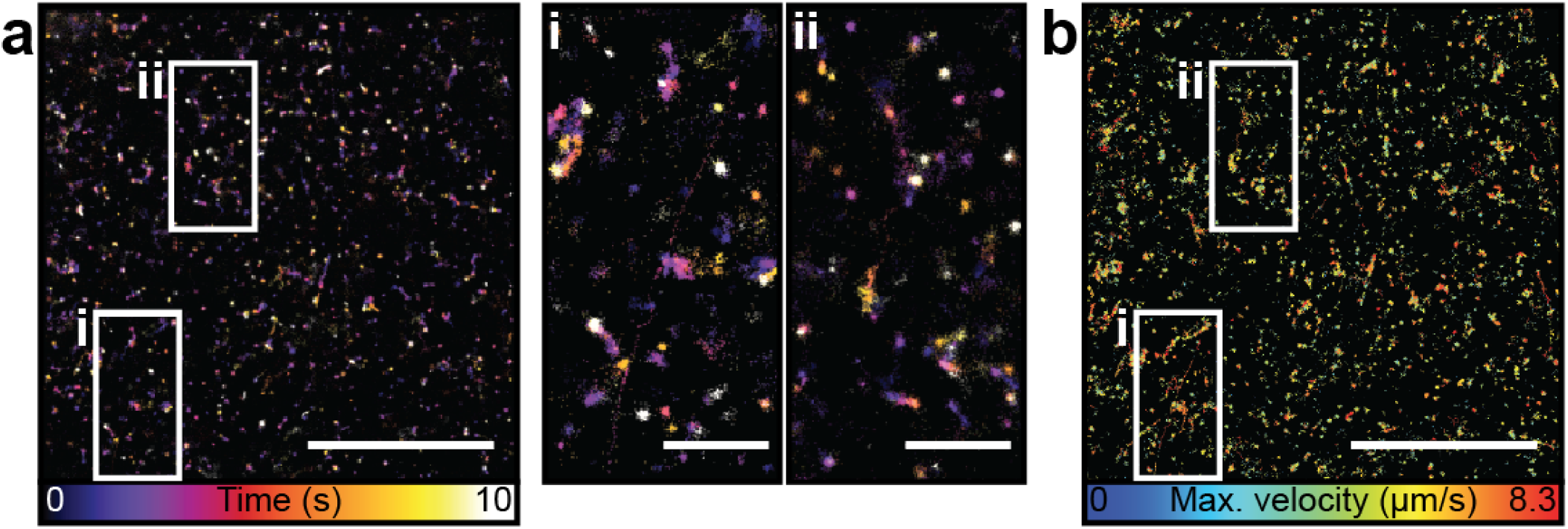
Live-cell SR single particle tracking with Super-Beacons. a) Histogram localizations from the first 10s of acquisition using Super-Beacon B3S9L, on 20 nm pixel grid coloured by time of localization. (I and II) Zoomed-in insets. Scale bar, 10 µm and 2 µm in insets. b) Single molecule tracks identified in a 30000 frame acquisition using the SB in live-cells. Tracks coloured by maximum velocity (highlighted regions are the same as in a). Scale bar, 10 µm.

Single-molecule tracking was performed using the single-particle tracking plugin for ImageJ, TrackMate (42) (Fig. 4b). Single molecules were identified in the raw images and detections were linked to form tracks using an algorithm based on the linear assignment problem proposed for single-molecule tracking in live-cells (42). Two frame gap closure with a maximum gap of 200 nm was performed. A total of 18667 tracks were detected, with a mean duration of 0.11 s and mean velocity of 2.97 µm s-1. The tracking results allow a single particle tracking image of the tracks to be generated (Fig. 4b). Consequently, SBs are a promising solution for the analysis of the diffusion of membrane-bound extracellular proteins on the molecular scale.

## Conclusion

Imaging of Super-Beacon labelled proteins in live-cells in low illumination regimes resulted in single-molecule localizations with precisions ∼ 30 nm and SMLM reconstructions with sub 100 nm resolutions, which demonstrates the potential of these probes in super-resolution. The ability to achieve this in a physiological buffer at low-illumination intensities demonstrates the merits of using SBs in live-cell settings. Furthermore, combining SB with techniques such single-particle tracking has the potential to allow experiment design using SB imaging to answer biologically relevant questions such as, in the example here, how IFITM proteins inhibit viral infection.

Limitations were identified, such as restrictions on labelling density due to steric hindrance and the biological relevance of ssDNA in live-cells. The size of the Ab conjugate hinders the use of SBs for live-cell imaging in their current form. The format used in this study, “Ab + streptavidin + SB”, entails a large distance between the probe and the target of interest. This translates in a large linker error visible in the cluster diameter measurements, for example (when compared with Ab directly conjugated to AF647). Other conjugations options are available if the reader wants to use SBs in live-cells. This option requires synthesising SBs with a Thiol modification instead of a biotin modification (also available from Metabion, Thiol-C6), this modification permits following a simple conjugation protocol already commonly used for DNA-PAINT (43). Alternatively, if the biotinylated SB solution seems more attractive due to the ease and cost-effect approach of using the same SB for both characterization and live-cell imaging, monomeric streptavidin is an option (44– 46). Additionally, reducing the illumination intensity to accommodate photo-toxicity and reducing the exposure time to accommodate dynamics both contribute a reduced number of detected photons. Finally, the duration of the on-states of the SBs presented are long in comparison to the optimal for fast imaging. Due to the nature of the transient quenching and the modularity and flexibility of DNA oligonucleotide design however, the possibilities for tailoring SBs to specific applications are extensive. For example, a 5bp stem can have 1024 different sequences. Together with different loop lengths and fluorophores there are thousands of possible 5bp stem SBs that could be considered. Additionally, the stability of the secondary hairpin structure can be modulated by the design of the oligonucleotide sequence, namely by altering the length of the hairpin and the avidity of the complementary stem region. The hybridization free energy can be determined by the identity of the bases in the stem sequence and experimental conditions such as pH, temperature or ion concentration and can be calculated using standard models of DNA hybridisation (47, 48). This stability can be further modulated in-situ by chemical environment and temperature. These factors combined open the possibility of designing SBs with bespoke properties for specific applications (e.g. imaging probes, as showcased here, or sensors) and/or environments (e.g. different salt concentrations or temperatures). Some of these properties such as temperature or chemical environment (49) can be changed dynamically at the microscope to further optimise their photoswitching behaviour while imaging. This flexibility suggests a model-based design principle could be adopted to reach an optimal structure for a specific application in order to maximally make use of the rapid imaging technologies available.

## ACKNOWLEDGEMENTS

This work was funded by grants from the UK Biotechnology and Biological Sciences Research Council (BB/M022374/1; BB/P027431/1; BB/R000697/1) (R.H., P.M.P.), the UK Medical Research Council (MR/K015826/1) (R.H.), the Wellcome Trust (203276/Z/16/Z) (R.H.). N.G. was supported by a PhD fellowship from the Engineering and Physical Sciences Research Council (EP/L504889/1). We would like to thank Dr. Siân Culley and Dr. David Albrecht (University College London) for critical reading and advice in the preparation of this manuscript.

## AUTHOR CONTRIBUTIONS

P.M.P, N.G. and R.H. devised the experiments. P.M.P. and N.G. acquired the experimental data sets and analyzed the data. M.M.M. and M.M. provided research advice and reagents. The paper was written by P.M.P., N.G. and R.H. with editing contributions of all the authors.

## Methods

### Super-Beacon probe design

The SB structure was designed to be short to reduce linker and crowding errors (1, 2) and to have a melting temperature above room temperature (∼ 33 C°-using mFOLD (3)) to favour the closed-state conformation (see characterization section). A SB was designed with the sequence 5’-ATTO550 ACG TTT T[Biotin-dT]T TTT CGT BHQ2-3’ called B3S9L-ATTO550-BHQ2, by the naming convention of having 3 base pairs in the Stem and 9 poly-T bases in the Loop, an ATTO550 fluorophore conjugated to the 5’ end and a BHQ2 quencher conjugated to the 3’ end (a control structure without the quencher was also designed 5’-ATTO550 ACG TTT T[Biotin-dT]T TTT CGT - 3’). Both structures were purchased pre synthesised, purified by high-performance liquid chromatography and lyophilized, from Metabion. Before use SBs were diluted to a concentration of 100 µM in TE buffer (10 mM Tris-HCL, 1 mM EDTA, pH 7.5) using the manufactures reported molar quantities and stored at −20°C, shielded from light sources. Both probes were designed with an internal modification to the thymine base at the center of the hairpin loop to a biotinylated thymine to allow the SBs to bind to streptavidin conjugated labelling agents such as antibodies (Ab) or be surface immobilized for *in vitro* characterization (Fig. 1d,e).

### Surface passivation for single-molecule studies

In order to determine the photoswitching characteristics of SBs, biotinylated SBs were immobilised on passivated surfaces. To do this 25 mm coverslips (HiQA - 1.5H, SAN 5025-03A, CellPath) were washed as previously described (4), followed by Piranha etching (5). Cleaned coverslips were amino-silanized and coated with NHS-ester polyethylene glycol (PEG-1-0001, ChemQuest Limited) and NHS-ester Biotin-PEG (PG2-BNNS-5k, ChemQuest Limited) as previously described (5). Coverslips were washed 10x with 1x liquid, sterile-filtered, Dulbecco’s Phosphate Buffered Saline plus 5mM MgCl_2_ (D8662, Sigma), filtered twice through a 0.22 µm filter and stored in a flask under permanent UV bleaching, from here on called “clean PBS”. Buffers were UV-bleached (and subsequently stores) under a 370 nm LED (370 nm LED, LedEngin Inc, LZ1-10UV00-0000), during all steps that are not affect by UV illumination (all steps prior to the addition of streptavidin to the surface). For this we used an open flask lid (e.g. 292270808, DWK Life Sciences), where the 370 nm LED attached to a heatsink (e.g. ICK LED R 23,5 × 14 G, Fischer Elektronik) was mounted.

Streptavidin was added to coated coverslips at 0.2 mgml^−1^ in 1x clean PBS for 10 min. Streptavidin coated coverslips were washed 10x with 1x clean PBS. SBs were diluted to a concentration of 10 nM to 100 nM and coverslips were incubated for 1 min to 10 min in order to have a low probability that multiple binding sites of streptavidin are occupied by a SB. After SBs were added, 100 nm TetraSpeck beads (T7279, Life Technologies), for drift correction, were added at a dilution of 1:1000 in 1x clean PBS for 10 min to each coverslip. Coverslips were again washed 10x with 1x clean PBS. On average number of detections in ‘empty’ coverslips prepared in an identical manner but without the addition of SBs was 0.090 ± 0.0092µm^−2^ at 561 nm illumination, this compares to an average of 0.38 ± 0.039µm^−2^ for ATTO550 SB coated coverslips indicating approximately 22% contribution of spurious noise detection from fluorescent debris in under 561 nm illumination.

### Illumination intensity determination

The reported illumination intensities are the peak intensities (*I*_peak_) at the centre of the field of view and were calculated from the percentage of maximum laser power provided by the microscope acquisition software, *p*. For each wavelength used, a conversion factor, *c*, was regularly calculated relating the software provided power to the measured power in the back aperture of the objective, *P* = *cp*. Over the course of the experiments a maximum of 10% error was measured. The transmission rate, *T*, of the objective was confirmed by measurement to be 89% ± 1.5%. For each wavelength used, the standard deviation, *σ*_*I*_, of the Gaussian distributed illumination intensity in the sample plane was measured by fitting the intensity of line profiles taken from illuminated empty coverslips. The calculation for *I*_peak_, assuming a Gaussian distributed illumination, *I*(*x, y*), of the sample, can be derived as follows, The maximum variation observed of the illumination intensity over the 5.2 µm imaged area was less than 5%.

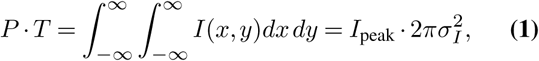

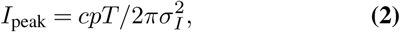

### Single-molecule imaging of Super-Beacon probes

Characterization was performed in phosphate buffer saline (PBS), using a range of illumination intensities, from live-cell compatible, 0.132 kW *cm*^−2^, to classical STORM intensities 2.64 kW *cm*^−2^ to determine the photoswitching properties of the SB probes. This was achieved through TIRF imaging of prepared coverslips using AttoFluor chambers (A7816, Thermo Fisher Scientific) in an ElyraPS.1 inverted microscope at room temperature (22°C). For excitation of ATTO550 a 561 nm laser operating at various illumination intensities was used. A 100x TIRF objective (Plan-APOCHROMAT 100x/1.46 Oil, Zeiss) was used, with additional 1.6x magnification, to collect fluorescence onto an EMCCD camera (iXon Ultra 897, Andor), yielding a pixel size of 100 nm. Imaging of ATTO550 SBs was performed with an exposure time of 18 ms using a cropped area of the camera (51.2 × 25.6 µm^2^) continuously for a period of 540 s (30000 frames).

### Single-molecule time-trace analysis

Data obtained was analyzed by extracting intensity over time traces (e.g. Fig. 1d) using a custom analysis pipeline as follows: Raw data was loaded with ImageJ/Fiji (6) and single molecule localization performed using the ThunderSTORM plugin (7). ThunderSTORM parameters were as follows: Filtering of the data was performed using a B-Spline Wavelet filter with scale 2, order 3. Detection of molecules was based on local maximum identification using 8 pixel neighbourhood connectivity. Detection threshold was set to 2 times the standard deviation of the first level of the wavelet decomposition. An integrated Gaussian PSF was used for localisation with initial *σ* = 160 nm. Fitting was performed using a weighted least squares estimate. Following localization of single molecules Thunderstorm was also used to calculate a per frame drift on the basis of fiducial marker localization’s using default settings. ThunderSTORM localization results, drift table and corresponding raw image data were loaded by a custom written MATLAB (Math-Works, Inc., USA) analysis program Results were filtered, retaining localisations with *σ*_*PSF*_ > 80 nm, *σ*_*PSF*_ < 200 nm, *σ*_*loc*_ < 40 nm and *I* > 40 photons, where *σ*_*PSF*_ is the fitted standard deviation of the Gaussian PSF, *σ*_*loc*_ is the ThunderSTORM calculated precision and *I* is the ThunderSTORM calculated photon count. This removes spurious fits to single pixels with narrow *σ*_*PSF*_ which are common in ThunderSTORM analysis, and low precision and low photon count localizations. The drift corrected, filtered localizations were subsequently grouped using a frame by frame search for localizations separated by less than 100 nm from the continuously updated mean position of the grouped localizations. Groups with mean positions separated by less than 100 nm following the full search were merged. Any groups of localizations with a number of members greater than 75% of the total number of frames were identified as fiducial markers and any groups within a radius of 2 µm of these fiducial markers were removed from the localization list. Localizations were clustered using DBSCAN (8) with a minimum number of cluster members, *minPts* = 2, and neighbourhood radius, *ϵ* = 80 nm (2x the maximum expected localization error after filtering). The cluster members were used to determine the average position of single molecules and any single molecules separated by less than 200 nm were discarded. The remaining molecule positions were used to extract 5 × 5 pixel (0.25 µm^2^) regions centred (without interpolation) on the molecule position from the drift corrected (without interpolation) raw data to produce a set of ‘trace images’. Any molecules that drifted into or out of the field of view during acquisition were discarded. The median of each full frame of the raw data was subtracted from the trace image frames. Due to the sparsity of the molecule detections this accounted for the contribution of illumination intensity variation to the background signal (approximately 2% across the time of acquisition). The background intensity of each frame of the trace images was calculated as the mean of the 16 boundary pixels and subtracted from the trace images. The photon conversion factor of the EMCCD camera was used to convert background subtracted camera counts in the trace images to photon number trace images. The photon number trace images were converted to photon number per frame traces by integration of the convolution of each trace image with a correctly normalized Gaussian PSF. The photon number per frame traces were subsequently converted to SNR per frame traces by division by the standard deviation of all boundary pixels in the convolved trace image sequence.

Denoising of the SNR traces was performed in an automated manner by thresholding of the detail coefficients of a wavelet decomposition of the trace (9–11) using the ‘wden’ function in MATLAB. Briefly, Wavelet decomposition of the 1D SNR trace was performed using a 6 level, Haar-Wavelet. Thresholds for each level are determined on a heuristic basis using Stein’s unbiased risk principle where the risk function has high SNR and a fixed form universal threshold 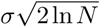 otherwise, where *N* is the dataset length (12). Thresholds are scaled by a level dependant estimation of the level noise, *σ*_*lev*_, in order to account for non-white noise. The detail coefficients are soft-thresholded and the denoised trace is computed from the wavelet reconstruction using the original approximation coefficients of level 6 and the thresholded detail coefficients of levels 1 to 6. The denoised SNR trace is hard thresholded at 10× the standard deviation of SNR values < 2 to determine on- or dark-state identities for each frame. Finally, transition times and state lifetime distributions were aggregated from the resulting idealized on/dark traces. The initial state was recorded separately in order to form initial on-state and initial dark-state distributions and the last state of each trace was discarded as corresponding to either an on-state truncated by the end of acquisition or an off state that is indistinguishable from a bleached state.

### Super-Beacon antibody conjugation

An anti-mouse secondary antibody (A21236 - Thermo Fisher Scientific) was conjugated to streptavidin using the Lightning-Link Streptavidin kit (708-0030, Innova Biosciences - following the manufacturer recommendations). After conjugation the Streptavidin-Antibody conjugate was incubated with 100 nM B3S9L-ATTO550-BHQ2-biotin SB or 100 nM B3S9L-ATTO550-biotin control for 16 h at 4C on a shaker in the dark. The resulting SB labelled Ab was purified via 100 kDa amicon spin filters (Merck - following the manufacturer recommendations).

### Super-Beacon for fixed cell imaging of microtubules

Fixed NIH3T3 cells were stained for β-tubulin (as previously described (13)) with SB (B3S9L-ATTO550-BHQ2), and control probe without quencher (B3S9L-ATTO550), conjugated secondary Abs. These samples were imaged with an illumination intensity of 0.132 kW *cm*^−2^ in PBS. The SMLM analysis of the SB labelled β-tubulin in fixed NIH3T3 cells data was performed using ThunderSTORM generating over 6×10^10^ localizations with a mean precision (*σ*_*l*_*oc*) of 22.0 nm (estimated by ThunderSTORM (7)). A low density region of the microtubule network was selected to judge the maximal achieved quality of the super-resolution image by calculating the full width half maximum (FWHM) on two representative line profiles perpendicular to microtubules. Because line profiles of microtubules can be narrowed by artefacts in localization algorithms (14), the limit of discrimination of the separation of two crossing microtubules was also measured.

### Super-Beacon Vs AF647 conjugated antibodies for fixed cell imaging of IFITM1

We imaged fixed cells (4%PFA at 23C for 30 min) stably expressing HA-tagged IFITM1 A549 cells stained with SBs (anti-HA Ab conjugated with streptavidin as previously mentioned) or AF647 (NHS ester conjugated) at increasing laser powers, from live-cell friendly 0.05 kW *cm*^−2^ to classical STORM illumination intensities 2.5 kW *cm*^−2^, in physiological PBS and toxic β-mercaptoethylamine (MEA), respectively. The SMLM analysis of the data was performed using ThunderSTORM (7) and to access the fidelity between the SMLM reconstructions and the raw data we used SQUIRREL ((15). To analyse IFITM1 distribution across a range of illumination intensities (from ∼ 0.05 kW *cm*^−2^ to ∼ 2.5 kW *cm*^−2^) we analysed the cluster diameter with SR-Tesseler (16).

### Super-Beacon SR single particle tracking of IFITM1 in live cells

A549 (IFITM1-HA) were incubated with SB labelled Anti-HA primary antibody for 30 min, at 37C in Ringers solution (cells were washed twice with Ringers solution before imaging). Live-cell imaging was performed at 0.132 kW *cm*^−2^, 561 nm illumination with exposure time 20 ms in Ringers. Single molecule tracking was performed using the single particle tracking plugin for ImageJ, Track-Mate (17). Single molecules were identified in the raw images and detections were linked to form tracks using an algorithm based on the linear assignment problem proposed for single molecule tracking in live-cells (17). Two frame gap closure with a maximum gap of 200 nm was performed. A total of 18667 tracks were detected, with mean duration of 0.11 s and mean velocity of 2.97 µm s-1.

## Supplementary Note 1: Supplementary Figures

**Fig. S1.**
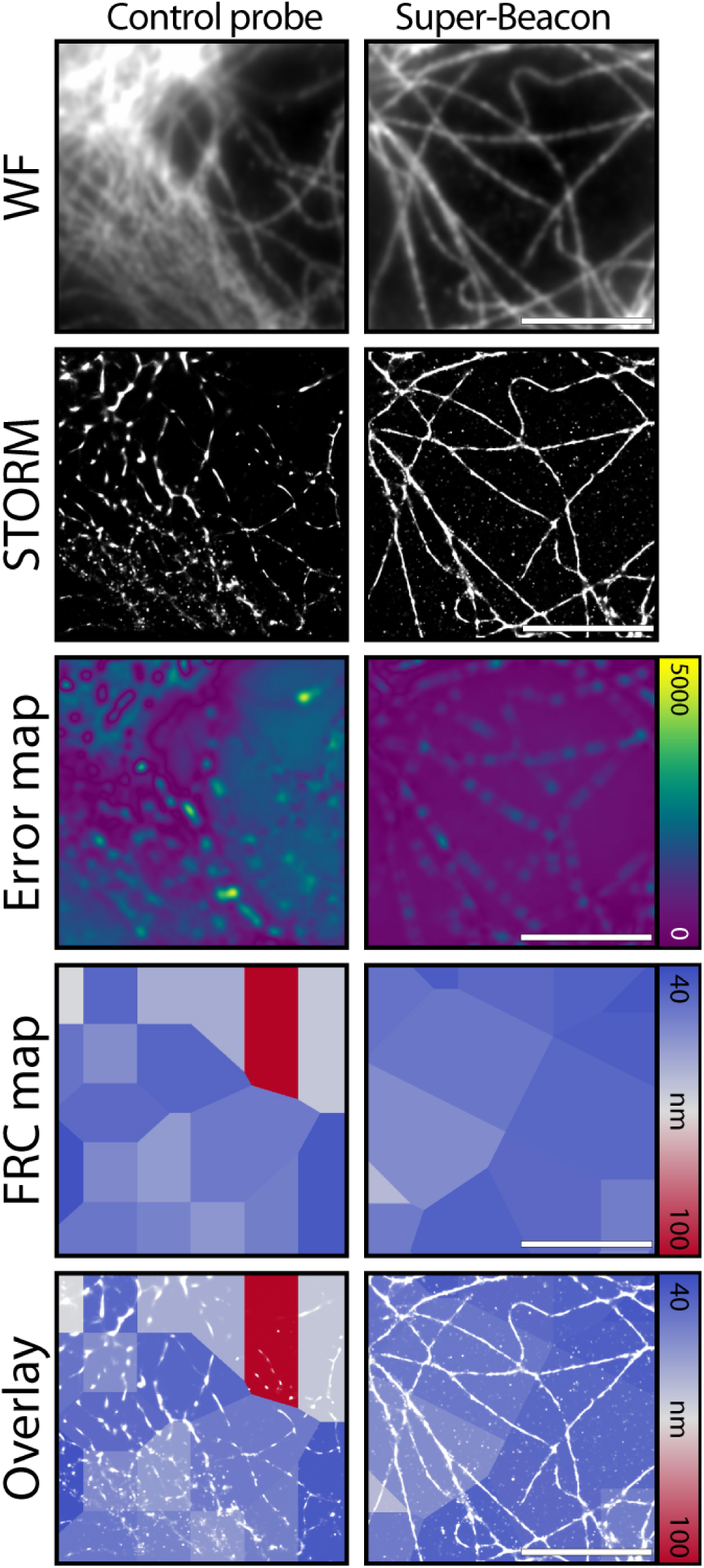
Fourier Ring Correlation (FRC) resolution mapping and NanoJ-SQUIRREL error maps of images acquired using SB or AF647 at different illumination regimes. a) Individual A549 IFITM1-HA expressing cells immunolabelled with anti-HA-streptavidin conjugated Ab labelled with B3S9L-ATTO550-BHQ2, WF snapshots (WF), SB super-resolution renderings (STORM), equivalent FRC map (FRC map), overlay between STORM and FRC map (Overlay), NanoJ-SQUIRREL error map of intensity normalized WF and STORM images. b) Same as in a) but for AF647 labelled Ab. Scale bars, 10 µm.

**Fig. S2.**
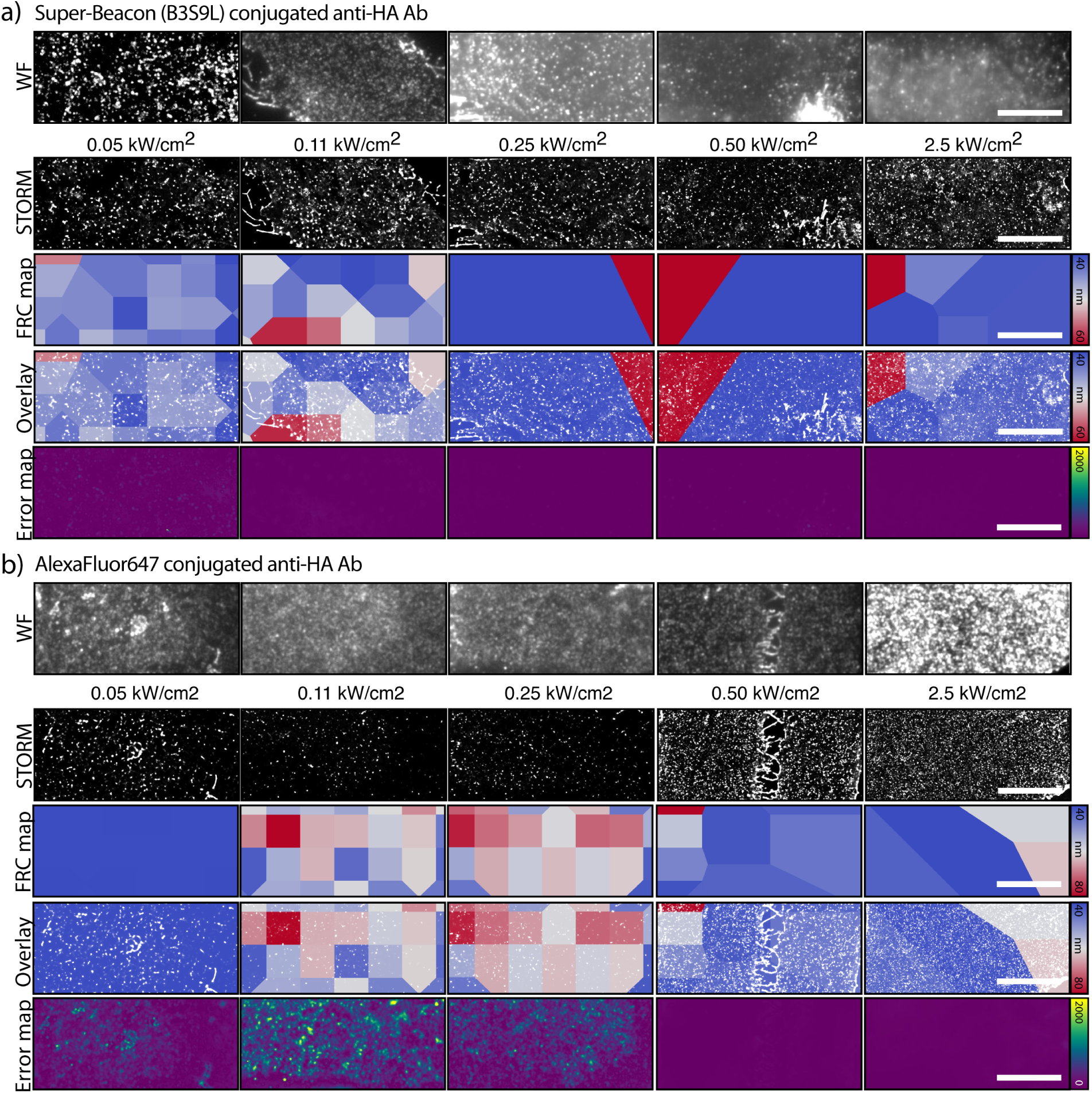
Fourier Ring Correlation (FRC) resolution mapping and NanoJ-SQUIRREL error maps of images acquired using SB or AF647 at different illumination regimes. a) Individual A549 IFITM1-HA expressing cells immunolabelled with anti-HA-streptavidin conjugated Ab labelled with B3S9L-ATTO550-BHQ2, WF snapshots (WF), SB super-resolution renderings (STORM), equivalent FRC map (FRC map), overlay between STORM and FRC map (Overlay), NanoJ-SQUIRREL error map of intensity normalized WF and STORM images. b) Same as in a) but for AF647 labelled Ab. Scale bars, 10 µm.

